# Fine-scale dynamics of functional connectivity in the face processing network during movie watching

**DOI:** 10.1101/2022.06.07.495088

**Authors:** Gidon Levakov, Olaf Sporns, Galia Avidan

## Abstract

Face are naturally dynamic, multimodal and embedded in rich social context. However, mapping the face processing network in the human brain and its relation to behavior is typically done during rest or using isolated, static face images. The use of such contrived stimuli might result in overlooking widespread cortical interactions obtained in response to naturalistic context and the temporal dynamics of these interactions. Here we examined large-scale cortical connectivity patterns measured in response to a dynamic movie in a sample of typical adults (n=517), to determine how inter-subject functional connectivity (ISFC) relates to face recognition scores. We found a positive correlation with recognition scores in edges connecting the occipital visual and anterior temporal regions and a negative correlation in edges connecting attentional dorsal, frontal default, and occipital visual regions. These ISFC patterns resembled previous findings comparing individuals with congenital prosopagnosia to normal controls and the viewing of inverted compared to upright faces. To further examine these connectivity patterns, we developed a novel method that allows analysis of inter-subject stimulus-evoked node/edge responses at a single TR resolution. Using this method, we demonstrated that co-fluctuations in face-selective edges observed here and in previous work are related to local activity in core face-selective regions. Finally, correlating this temporal decomposition of the observed ISFC patterns to the movie content revealed that they peak during boundaries between movie segments rather than during the presence of faces in the movie. Our novel approach demonstrates how visual processing of faces is linked to fine-scale dynamics in attentional, memory, and perceptual neural circuitry.

## 1. Introduction

Face recognition is a highly developed visual skill that is pivotal for human social interactions. Face processing requires the integration of multiple brain regions such as the fusiform face area (FFA), occipital face area (OFA), and the superior temporal sulcus (STS) that respond selectively to faces compared to other visual categories^1^. Outside these ‘core’ face-selective patches, additional ’extended’ regions^2–4^ are implicated in the processing of memory and identity (anterior temporal lobe; ATL^5, 6^), attentional (intraparietal sulcus; IPS^7^), and emotional (amygdala^8^) aspects of faces. Interactions among these ‘extended’ regions are typically observed in response to naturalistic audiovisual face stimuli that are dynamic and embedded in rich social context^3, 9, 10^. However, little is known of how these interactions and their dynamics are related to face recognition abilities

Functional connectivity studies have revealed the importance of interactions among core and extended face regions to the successful recognition of faces. For example, two studies examined the effect of disrupted face processing on functional connectivity and showed that face inversion^11^ or a congenital impairment in face processing (congenital prosopagnosia; CP^12^) were associated with hyper-connectivity in occipital and lateral visual regions. In contrast, intact face processing was linked to connectivity to medial and anterior portions of the ventral visual pathway. While these studies used static face images, individual differences in face recognition in the general population^13, 14^ have only been tested during rest. Moreover, these studies were limited to testing connectivity within^15, 16^ and from^17^ ’core’ face-selective regions, overlooking interactions in ‘extended’ regions. Thus, widespread interaction among ‘extended’ regions and their response to naturalistic face processing remains largely unexplored.

The usage of movies in neuroimaging allows us to measure subjects’ shared functional interactions in response to rich, social naturalistic stimuli^18^. Inter-subject functional connectivity (ISFC) is a computational method that measures these interactions between the brains of different individuals driven by a common stimulus^19^. Current methods for quantifying frame-wise co-fluctuations in functional connectivity^20^ allow for assessing brain responses in both rest and movie data at fast temporal scales without the need for windowing. Here we present a novel method combining these two approaches to decompose ISFC values to the single time frame temporal resolution. We term this approach the inter-subject node/edge time series (IS-NTS/ETS) and demonstrate how it decomposes connectivity-behavior correlations to their fine-scale spatial and temporal dynamics.

The present study aimed to test how individual differences in face recognition relate to dynamic functional connectivity patterns observed as subjects watch a movie rich in social interactions. Specifically, we focused on the following questions (Fig. 1): 1. **What** cortical-wide interactions correlate with face recognition scores in the general population? 2. **Where** in the brain are such functional interactions related to local activations? and 3. **When** do these functional interactions occur in time with respect to the content of the movie? To address these questions, we utilized a large adult lifespan data set (N = 517; Cam-CAN^21, 22^) containing functional magnetic resonance imaging (fMRI) data acquired during movie watching and behavioral face recognition scores. We divided participants into groups according to their face recognition scores while keeping age distributions equal. We then correlated their ISFC values to the behavioral group affiliation. We tested the relation between brain-wide functional interactions and local activations using a novel method we termed ‘edge seed correlation’. Finally, using data from an independent group of human raters^23^ and a deep learning face recognition algorithm^24^, we examined which stimuli in the movie preceded the emergence of the IS-ETS correlations with behavior.

**Fig. 1.**
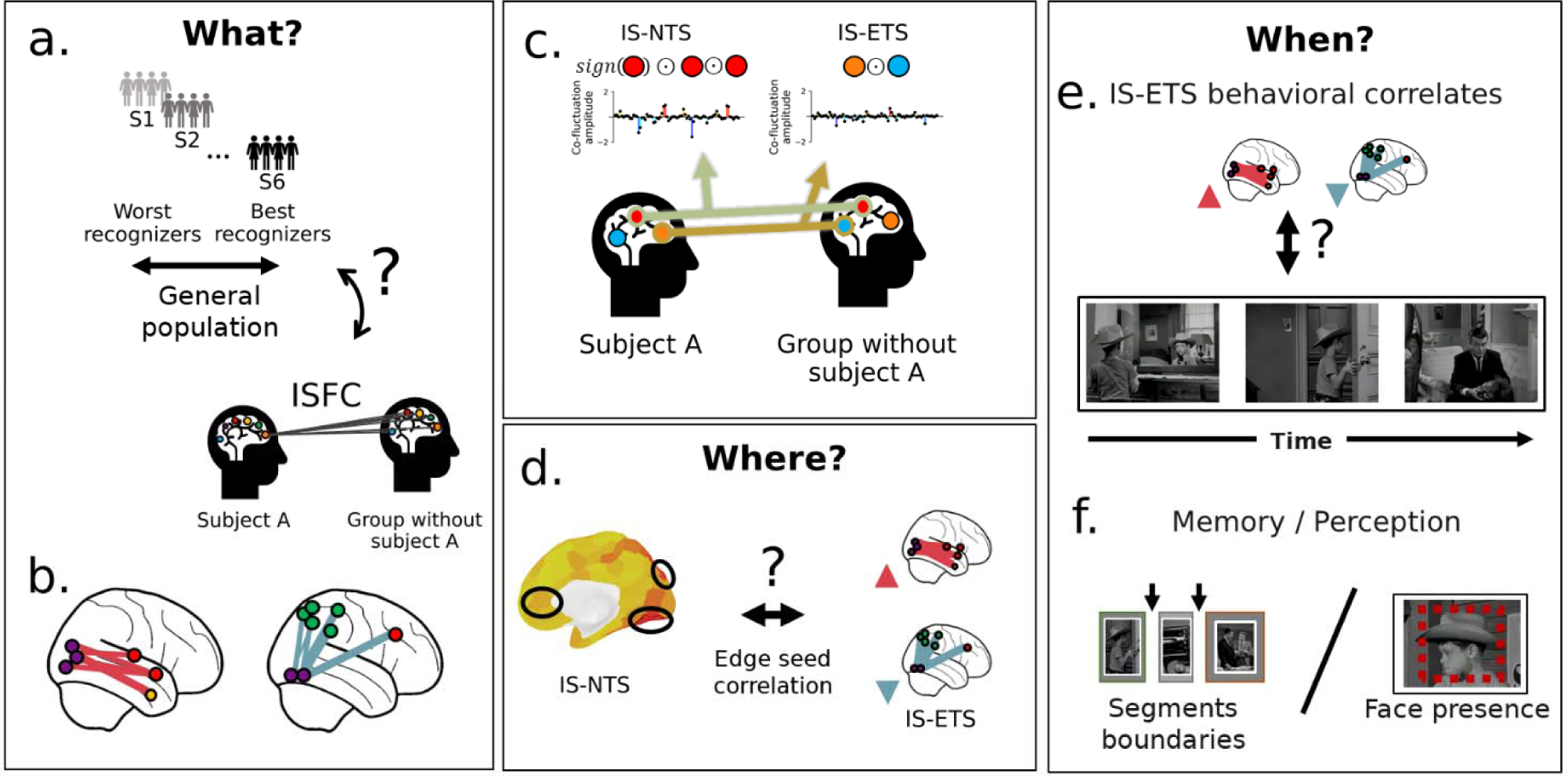
Study outline and outstanding questions. What - (a) We tested what are the ISFC correlations of face recognition in the normal population. (b) This test results in a set of edges positively (red) or negatively (blue) correlated with face recognition scores. (c) We present a novel method that decomposes the inter-subject connectivity among homologous (e.g., same voxel or brain region) or different brain regions in time to their edge time series ^20^. When applied to homologous brain regions (inter-subject node time-series; IS-NTS) is computed as the element-wise multiplications of the z-scored signals of two nodes across subjects. The time series is then multiplied by the sign of the nodes’ signal to differentiate co-activations and co-deactivations. When applied across different brain regions, (inter-subject edge time series; IS-ETS), the same procedure is used without the last multiplication step. (d) In a procedure we term ’IS-edge seed correlation’, the two measures are correlated to reveal whether local brain activations (seed) may recapitulate the cortical-wide correlations of successful face recognition. (e) We then examine when, during the movie, these connectivity-behavior correlations appear. (f) We tested whether these neural-behavioral correlates would be related to the visual perception of faces or to their encoding in memory.

## 2. Results

### 3.1 ISFC differences and face recognition behavioral performance

Previous work examining functional connectivity correlates of face recognition was conducted during rest, while faces are naturally dynamic, multimodal, and engaged in social interactions. Here we tested the correlation between participants’ ISFC, taken while watching a socially enriched movie, and face recognition scores. We quantified face recognition as the first PC of Benton’s face recognition test and a famous faces test which accounted for 67.57% of the two variables’ variance. Participants were divided into six groups according to their combined face recognition score while keeping age distribution equal across groups (Table 1; see methods section 2.7). We found that dividing the subjects into six groups optimally balanced sample size (n= 84-90) and within-group behavior homogeneity. We replicated all results using different choices of the number of sub-groups (*i*=5,7; SI 9.1).

**Table 1.** Demographics and face recognition scores for each behavioral group.

| Group | Age | Handedness | Gender | Benton | Famous | Combined face score | N |
| --- | --- | --- | --- | --- | --- | --- | --- |
| **S1 (worst)** | 52.2<br>($\pm 18.6$) | 78.7<br>($\pm 44.7$) | F=49, M=41 | 0.77<br>($\pm 0.07$) | 0.57<br>($\pm 0.11$) | -1.38<br>($\pm 0.71$) | 90 |
| **S2** | 52.1<br>($\pm 17.5$) | 65.1<br>($\pm 65.4$) | F=38, M=46 | 0.81<br>($\pm 0.07$) | 0.66<br>($\pm 0.08$) | -0.45<br>($\pm 0.67$) | 84 |
| **S3** | 52.5<br>($\pm 17.6$) | 82.2<br>($\pm 42.6$) | F=44, M=42 | 0.84<br>($\pm 0.07$) | 0.69<br>($\pm 0.06$) | 0.03<br>($\pm 0.64$) | 86 |
| **S4** | 52.4<br>($\pm 17.3$) | 76.8<br>($\pm 54.5$) | F=37, M=47 | 0.87<br>($\pm 0.06$) | 0.71<br>($\pm 0.05$) | 0.38<br>($\pm 0.56$) | 84 |
| **S5** | 51.5<br>($\pm 17.6$) | 89.4<br>($\pm 32.6$) | F=45, M=39 | 0.90<br>($\pm 0.05$) | 0.73<br>($\pm 0.04$) | 0.75<br>($\pm 0.49$) | 84 |
| **S6 (Best)** | 52.8<br>($\pm 16.8$) | 77.4<br>($\pm 53.1$) | F=48, M=41 | 0.95<br>($\pm 0.04$) | 0.74<br>($\pm 0.02$) | 1.25<br>($\pm 0.41$) | 89 |
| **P<sub>(difference)</sub>** | 0.998 | 0.059 | 0.718 | $< 2.2e-16$ | $< 2.2e-16$ | $< 2.2e-16$ | |

We measured Spearman’s rank correlation between the behavioral group affiliation and ISFC values. We compared these to a null distribution generated by shuffling the group affiliations, recomputing the ISFC values and the correlation to behavior. We considered an edge significant if its correlation values were higher/lower than the entire null distribution (*k=10,000*). Fig. 2 depicts a glass brain projection of the positively (‘positive configuration’) and negatively (‘negative configuration’) correlated edges to behavior along with results using a more lenient threshold (p<0.001). We found that edges that negatively correlated with recognition scores were mostly between visual and attentional dorsal (13.04%) or frontal default (60.87%) network nodes. The dorsal attentional nodes are part of the bilateral posterior parietal cortex (PPC), and the frontal default nodes are mainly composed of the medial prefrontal cortex (mPFC) and the inferior frontal gyrus (IFG). Only three positively correlated edges were found significant. Hence, we additionally examined edges using a more lenient threshold (p<0.001). This revealed that the nodes with the highest degree for the negative configuration were the rh. lateral occipital cortex (LOC) and the mPFC (both degree=14). The lh. ATL was the highest degree node for the positive configuration (degree=14). This pattern of hyper-connectivity to occipital visual regions for the negative configuration and hyper-connectivity to the anterior temporal region for the positive configuration was previously found when comparing intact to disrupted face processing^11, 12^. Hence, we examined whether these results may be driven by the group with the lowest face recognition scores. To do so, we tested the change of the correlation to behavior within the highest correlated edges (p<0.001 threshold) following the exclusion of each group. Excluding the group with the lowest scores (S1) did not result in a smaller effect than removing each intermediate group (S2-S5; t=-0.068, p=0.946). We also did not find such an effect when excluding the group with the highest scores (S6; t=0.079, p=0.937; see SI Fig. 5). These results imply that ISFC correlates of face recognition spread to multiple regions outside the ‘core’ face regions. These findings resemble connectivity patterns found during a disruption of typical face processing.

**Fig. 2.**
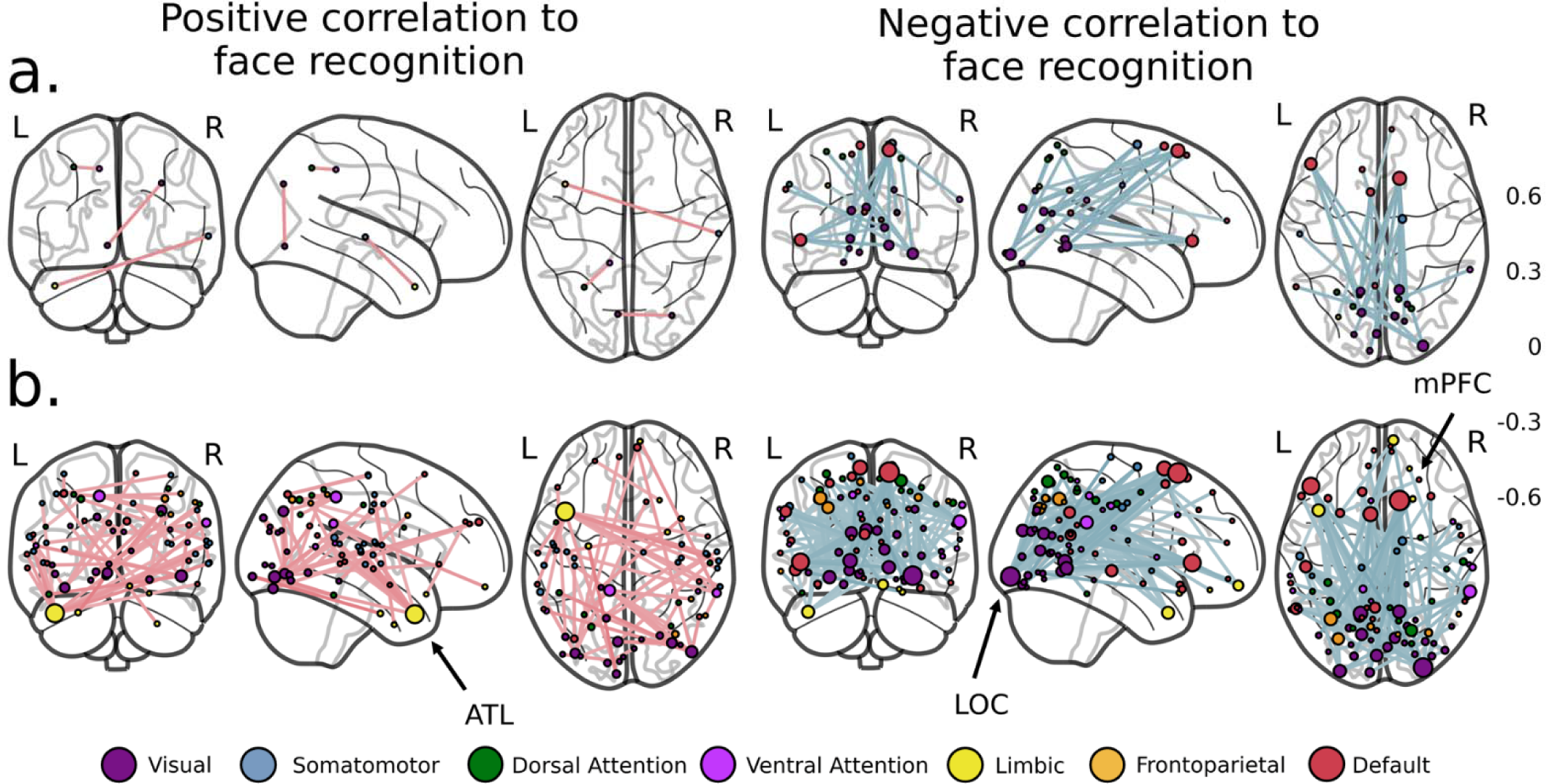
Correlation of ISFC values and face recognition ability. We computed Spearman’s rank correlation of ISFC values and the behavioral group affiliation for all edges. We compared the correlation coefficients to a null distribution generated by shuffling the group affiliations, recomputing the ISFC values and the correlation to behavior. Results are depicted on a glass brain separately for positively (left) and negatively (right) correlated edges. On both panels, the upper (a) represents edges that were larger (left) or smaller (right) than the entire null distribution (k=10,000). We include results using a more lenient threshold (p<0.001) on the lower panels (b). The left anterior temporal cortex (ATL) was found as the node with the largest degree in the positive network (degree=14) and the left LOC and mPFC in the negative network (both degree=14). The color of the edges indicates their correlation to behavior. The color of the nodes indicates their affiliation to one of the seven canonical resting-state networks ^25^. Nodes’ size is proportional to their strength, i.e., the sum of their absolute edge weights. Edges were projected onto coronal (left), sagittal (middle), and horizontal (right) views.

### 3.3 Neural modulation of the optimal and sub-optimal configurations of the face network

We asked whether the observed network of positive and negative edge correlation to behavior (see 3.1) and similar findings observed during disrupted face processing^11, 12^, could relate to activations in specific brain regions. To answer this, we applied our novel IS edge seed correlation method (see 2.9) that correlates the activity of a given brain region (seed) to stimulus-driven co-fluctuations in all possible edges in the brain. To validate the approach, in SI section 9.4, we include an application of this method to the frontal insula and replicate previous findings regarding its role in switching between the default mode and the frontoparietal networks^26, 27^. We term the patterns of successful and disrupted face processing found here and in our previous work, as the optimal and sub-optimal face processing configurations. We defined the sub-optimal configuration as connectivity within the visual, dorsal attention, and default-A networks. We defined the optimal configuration as connectivity within the visual network and the anterior portion of the temporal lobe defined as the ATL and the tempo-parietal and limbic-A networks. We compared the dice similarity of the optimal and sub-optimal configurations to the positive and negative IS edge-seed correlations of all brain regions, respectively. We plotted the results of the mean dice similarity over the two configurations on the brain surface (Fig 3a) along with the edge-seed correlation of the rh. posterior FFA (pFFA) in which we found the highest similarity (Dice=0.092; Fig. 3b). Comparing the effect size gained with the IS (Mean Cohen’s d: 4.88) versus within-subjects version (Mean Cohen’s d: 2.72) of the edge seed correlation method revealed a larger effect when using the IS variant (t(1599)=191.464, p<2.2e-16). The spatial arrangement of the similarity values visually resembled the core face-processing network, suggesting that the appearance of the two configurations is uniquely related to co-activations and deactivations within face-selective regions. We decoded our cortical map to quantify this potential similarity by matching it to a dictionary of 128 cognitive brain maps (Cogspaces^28^). We found that the map most strongly represented by the word “face” had the highest dice similarity to our map (Fig. 3c; Methods 5.11). We asked whether these results are specific to face-selective regions or observed in other high-level visual areas. Hence, we conducted a similar analysis with the bilateral parahippocampal place area (PPA), a high-level visual area spatially and functionally distinct from the FFA^29^. The results did not show a similar effect (Dice: rh. PPA=0.011, lh. PPA=0.007; SI Fig.6). These findings imply that co-activations and deactivations in face-selective regions are uniquely associated with the appearance of the two optimal and sub-optimal face processing configurations.

**Fig. 3.**
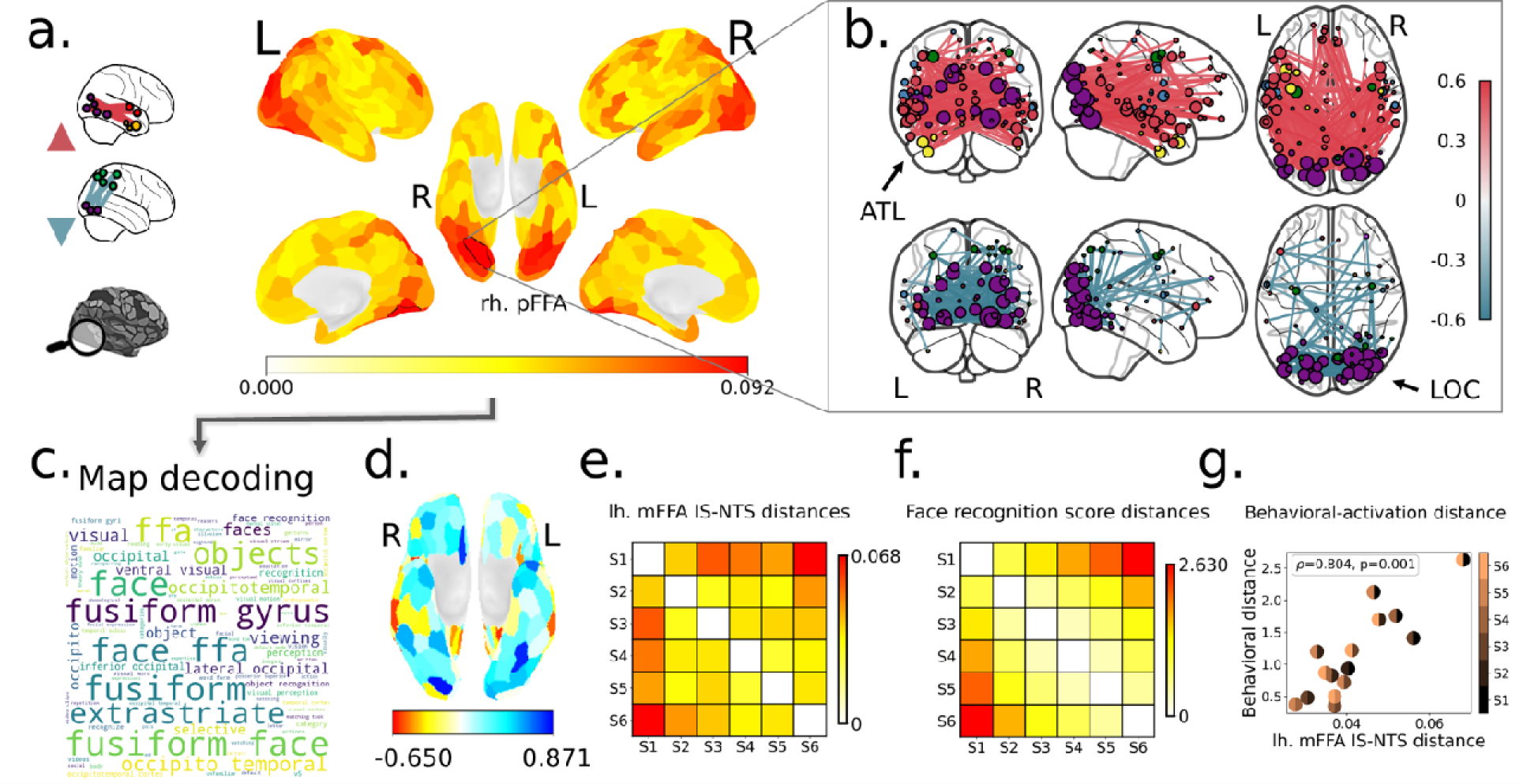
Brain regions synchronized with optimal and sub-optimal face network configurations. (a) Surface plot of the dice similarity between each brain region IS edge seed correlation and the optimal and sub-optimal configurations. (b) Results of an IS edge seed correlation of the rh. pFFA (p<1.0e-05), in which we found the highest similarity (Dice=0.092). (c) Word cloud visualization of the closest cognitive brain map chosen from 128 possible maps (Cogspaces dictionary; Mensch et al., 2021). (d) Ventral view surface plot depicting the correlation of regional activation differences to groups’ behavioral differences. Matrices representing the cross-group differences in behavior (e) and activation (f) and a scatter plot of the two (g) are shown for the lh. mFFA, in which we found a significant correlation.

### 3.3 FFA activity and individual differences in face recognition

We hypothesized that given the role of the FFA in face processing^30^ and its correlation to the appearance of the two configurations during the movie, its signal would relate to face recognition scores. Thus, we tested whether differences in the FFA’s IS-NTS across groups would be correlated with group differences in behavior. We calculated differences in behavior using a Euclidean distance of the face recognition score and activation differences as one minus the correlation of the FFA IS-NTS across groups. We found a significant correlation in the lh. medial FFA (mFFA; ρ=0.804, p=0.001; Fig. 3e-g) while a similar trend in the lh. pFFA that did not reach statistical significance (ρ=0.521, p=0.055; see SI Fig.7 for whole-brain analysis). We did not find the same correlation in the right hemisphere (both ρ’s <0.232, p’s>0.254).

These results suggest that face recognition is related to a distinct activation pattern within the lh. FFA in response to a common stimulus that contained faces. We discuss possible reasons for the left laterality found here and within the left ATL (see 3.1) in the discussion.

### 3.4 Temporal predictors associated with memory versus perception and their relation to ISFC differences in face recognition

To better understand the cognitive functions associated with ISFC differences in face recognition, we took “a reverse engineering” approach and probed which stimuli appeared in the movie before the emergence of the correlations to behavior. The IS-ETS framework allows us to probe such questions as they decompose the ISFC to a temporal resolution of a single time frame. One straightforward prediction is that these patterns are related to the processing of visual aspects associated with the faces. To test this hypothesis, we extracted a set of face perception predictors from the movie using a pre-trained deep face recognition model. Each frame contained between zero and five faces and overall faces appeared in 71.0% of the frames (see Methods 5.5 for the code and all extracted measures). The predictors included the presence of faces, social interactions depicted in a scene (i.e., two faces or more within a scene), the number of faces in the scene, and faces’ size within the scene (Fig. 4c). A second hypothesis was that these patterns of connectivity were related to the encoding of faces in memory. Previous work suggested that the encoding and binding of perceived information in memory occur during boundaries between movie segments^31, 32^. Accordingly, we used these movie boundaries as memory-related predictors, rated by independent observers in a previous study^23^. ’Neural events’, brain-wide co-fluctuations were additionally used as neural predictors due to their substantial contribution to functional connectivity variance and their response to naturalistic stimuli^20^ (Fig. 4b). To quantify the emergence of the positive and negative configurations, we derived the IS-ETS of every subject for each time frame and computed their correlation with the behavior group affiliation. The resulting edge correlation maps was compared to the global edge correlation map obtained using the ISFC (Results, section 3.1) using dice similarity. We repeated this for positive and negative correlations to behavior. As the two positive and negative similarity time series were highly correlated (r=0.479, p=6.6e-12), in all following analyses, we took the average of the two dice coefficients (‘mean similarity’). We convolved all movie-related predictors with the HRF to account for the hemodynamic delay. We tested the correlation between all possible pairs of the four face-related predictors (six comparisons). All face-related predictors were positively correlated among themselves (all r’s>0.207, all p’s <0.036) except for the correlation between the number of faces and face size (r=0.157, p=0.081). There was a small, marginally significant correlation between movie boundaries and neural event measures (r=0.136, p=0.049). Importantly, both perceptual movie boundaries (r=0.148, p=0.004) and neural events (r= 0.293, p<1.0e-04) correlated with the emergence of the global configurations while none of the facial predictors reached significance (all r’s<-0.026, all p’s>0.650; Fig. 4d). We additionally analyzed movie boundaries as discrete rather than continuous. We tested the mean similarity to the global configuration following each boundary, using a 2 TR lag (∼5s) accounting for the hemodynamic delay^23^. The similarity to the global pattern was significantly larger than a null distribution derived by randomly shifting the boundaries in time (p=0.016; Fig. 4e). Although the face perception predictors are more continuous in nature, we conducted a similar analysis as a control in a matched number of frames were the face stimulus size or the number of faces in a scene were maximal. Both stimulus size (p=0.786; k=10,000) and the number of faces (p=0.505; k=10,000) did not reveal similar findings.

**Fig. 4.**
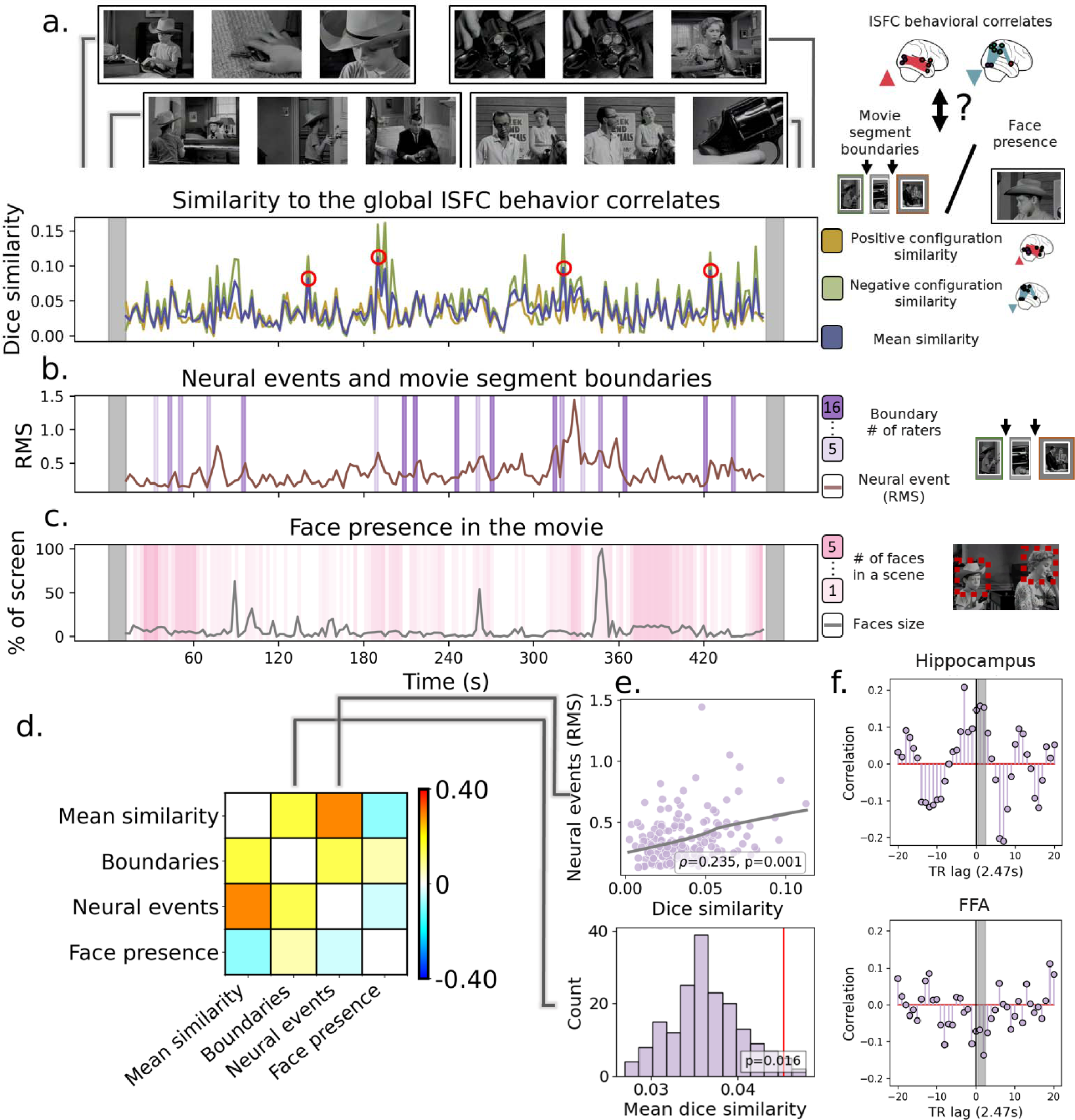
The relation of memory and perception to the emergence of positive and negative configurations. (a) We derived the IS-ETS of all subjects and computed its correlation with the behavioral group affiliation in each time frame. The resulting time series (bottom) indicates which points in time exhibited high dice similarity with the positive and negative configurations and their mean. The top part of the figure depicts the movie frames that preceded the peaks in the mean similarity time series (-2 TRs, ∼ 5s) and frames that appeared ∼10s (4 TRs) before (left) and after (right) this frame to provide its context. (b) A plot of neural events^20^ along with a marker of frames that followed perceptual movie boundaries shifted +2 TR (∼5) to account for the hemodynamic lag. The color of the windows indicates the number of observers that rated them. (c) Plot of face size presented at each movie frame along with windows following the appearance of faces on the screen (shift forward in time as in b). Windows colors indicate the number of faces in the scene. The shaded region in the time series (a,b,c) marks excluded time frames at the beginning and the end (5 TRs; ∼12s). (d) Correlation matrix depicting the correlation between the mean similarity time series and movie segment boundaries, neural events and face presence. (e; top) Scatter plot of the relation between neural events and the mean similarity. (bottom) A histogram depicting the mean similarity to the global configuration following movie segment boundaries (red line) compared to a null distribution created by shuffling the boundaries in time. (f) The correlation between the mean similarity and the IS-NTS of the hippocampus (top) and the FFA (bottom). We present the correlation as a function of the delay in time.

Next, we asked whether activity within brain regions related to the visual processing of faces and the encoding in memory of movie segments may provide additional support to one of the two hypotheses. We used the mean activity of the posterior and medial FFA, known for its involvement in the visual perception of faces^33^, as a neural marker of face perception. Similarly, we used the hippocampus as a neural memory encoding marker due to its role in the encoding of perceptual segments into memory^34^.

Previous work reported delayed activity in the hippocampus that was longer than the typical hemodynamic lag (>6s^32^). Hence, we also tested the delayed correlation of both regions (lag=0,1,2 TRs). We used the mean IS-NTS of the right and left hippocampus as we did not have an *a priori* hypothesis regarding a laterality effect. The hippocampus presented an immediate (r=0.146, p=0.036) and delayed (+1 TR: r=0.157, p=0.029; +2 TRs: r=0.153, p=0.032) correlation to the global configuration emergence. A similar delayed or immediate correlation in the FFA did not reveal a similar pattern (all r’s r<-0.068, p>0.839; Fig. 4f) nor when analyzing the right and left FFA separately (all r’s r<-0.038, p>0.711). Finally, we conduct a post-hoc analysis to further examine the memory encoding hypothesis. We suggest that hippocampal-FFA connectivity supporting reactivation of face representations^35^, might correlate to face-recognition scores. A correlation to recognition scores for the lh. pFFA-hippocampus connectivity did not survive a correction for multiple comparisons (ρ=0.150, p=0.035, p_FDR_=0.140; SI Fig. 8). Similar analysis within the lh. mFFA or the rh. m/pFFA (all ρ’s<0.115, p’s>0.091) were not significant.

## 4. Discussion

Face recognition is a complex cognitive ability that depends on integrating perceptual, attentional, and memory neural circuits. The current work examined how dynamics in these cortical-wide interactions, evoked by an audiovisual stimulus that contained faces, related to face recognition scores. We found that correlations of cortical-wide ISFC to recognition scores were widespread and observed outside the ‘core’ face-selective regions. Interestingly, these patterns resembled previous findings observed in studies comparing individuals with congenital prosopagnosia to typicals^12^ and the viewing of inverted compared to upright faces which hampers face processing among typical observers^11^. We present a novel methodology that assesses fine-scale temporal dynamics in inter-subject functional connectivity. Using this method, we showed that ISFC patterns could be related to activations in the core face-selective regions. Finally, decomposing the ISFC correlations to behavior in time revealed that they are likely associated with encoding faces in memory rather than with their perception.

### 4.1 Connectivity patterns related to sub-optimal processing of faces

Our primary analysis (see 3.1) outlined the set of cortical interactions related to face recognition scores in the normal population. Overall, negative correlations to behavior included connections between occipital visual regions, mainly the LOC, and the PPC, the mPFC, and the IFG. The PPC is involved in processing observed actions^36, 37^ and modulating attention during visual processing^38, 39^. Specifically, evidence suggests that the PPC modulates functional connectivity between early visual cortex and high-level category-specific visual regions in response to increased task demands^40^. In line with these findings, transcranial magnetic stimulation (TMS) to the bilateral PPC was shown to disrupt configural but not featural processing of faces^41^. In addition to the PPC, we observed multiple visual connections to the mPFC and the IFG, regions implicated in the dynamic processing of faces^42–44^ and the processing of familiar faces^45, 46^. In the context of connectivity rather than regional activations, functional connectivity between visual^47^ or temporal^48^ cortices and the prefrontal cortex is related to the processing of degraded or ambiguous faces or social visual stimuli. Moreover, differences in activations of these frontal face regions were found in Autism Spectrum Disorder (ASD) compared to control while observing faces^49, 50^. Hence, these occipital-frontal^47^ and occipital-parietal^51^ connections may represent a top-down compensatory mechanism in response to sub-optimal processing of faces. Such a hypothesis may explain similar patterns found while observing inverted compared to upright faces^11^.

### 4.2 Connectivity pattern related to optimal processing of faces

In contrast to the negative correlations, positive ISFC correlations to behavior were less apparent, with only three significant edges. One of these edges connected the ATL, which was also found as the node with the largest degree when reducing the significance threshold. Neuroimaging^5, 52^ and neurological^53^ studies demonstrated that the ATL relates semantic knowledge to a perceived familiar face. Also, reduced structural^54^ and functional^12^ connectivity was found between ventral face regions and the ATL in CPs compared to controls. Most studies reported that disrupted face processing is related to reduced connectivity between ventral visual areas such as the FFA and ATL^55^. In contrast, we found most connections to the ATL in regions along the STS. We hypothesize that this is due to the usage of dynamic stimuli that included motion, audio, and body images compared to static images used in previous studies. Responses to dynamic^3, 9^, audiovisual^10^ face stimuli are typically more pronounced along the STS. Additionally, viewing movies with faces that included speech^56, 57^ social interactions^58^ and bodies^59^ also showed a leftward bias in line with the known left lateralization of language perception^60^. These findings may account for the left rather than right ATL hyper-connectivity found in the present study in contrast to previous results^12^.

### 4.3 Similarities of the results with cases of face processing impairments

Both negative and positive correlations with behavior resembled previous work comparing individuals with Congenital Prosopagnosia (CP) to typicals^12^ and the viewing of inverted compared to upright faces^11^. For example, in the current study, as well as in both previous studies, we observed hyper-connectivity in occipital visual regions and an increase in the connectivity of these regions to the PPC in the context of sub-optimal processing of faces. In contrast, optimal face processing, associated with increased connectivity to the ATL was found in the current study and also when comparing CP to typicals. These similarities emerge despite profound methodological differences such as separate samples, sites, clinical versus typical populations, preprocessing strategies, functional connectivity measures, and nodes definition. This can be viewed as evidence supporting the generalization of the results and imply that the three conditions may share common neural representations.

### 4.4 Discrete versus continuum of face recognition abilities

Face processing abilities vary considerably within the normal population. Studies have focused on two extreme cases, individuals with exceptionally high face recognition abilities (super-recognizers)^61^ on the one hand and individuals with CP, in which face processing is severely impaired on the other hand^62^. A critical question that emerges concerning these populations, which is still unresolved, is whether these cases fall within the tails of the normal distribution or whether they belong to a distinct population^63^. Such issues also arise in other disorders such as ASD^64^ and, while being very critical, are notoriously difficult to resolve. Here we found that negative ISFC correlations to behavior in the normal population resembled findings observed in individuals with CP^12, 65^. Moreover, the results were not driven by individuals with the highest/lowest recognition scores. Future studies that will test typicals and CPs using a similar experimental design, would allow to better pinpoint whether CPs indeed present qualitatively similar connectivity patterns. Such a design may provide more direct evidence that would shed light on whether CP entails a distinct population or rather the tail of the normal distribution. By the same token, characterizing connectivity in individuals with superior face recognition skills^66^ using such a design may enable us to explore the same issue but focusing on the other extreme end of the face recognition distribution.

### 4.5 Neuronal modulation of the optimal and sub-optimal face processing configurations

In neuroscience, we often explore how behavior correlates with functional connections^67^. Similarly, we can ask how the signal in a given brain region (‘seed’) may covary with, or modulate functional connectivity between two other brain regions (‘targets’). Such high-order interactions may stem from direct or indirect connections between the ‘seed’ region and one or both ‘targets’. Previous work explored such high-order interactions^68^ and a modulatory effect of specific nodes on brain-wide connections has been reported^26, 27^ (see SI 9.3 for our replication of these latter studies). However, here we present an innovative perspective for exploring these interactions in a stimulus-driven manner. We propose a novel method correlating the IS-NTS of a specific region to brain-wide IS-ETS. Using this method, we found that positive edge seed correlations of core face-selective regions were similar to the optimal face processing configuration. Likewise, negative edge seed correlations resembled the sub-optimal configuration. We can view these core face regions as high-order hubs for optimal and sub-optimal face processing networks. Moreover, group distances between behavior and the activations of these regions (see 3.3) were correlated, suggesting that the activation patterns of these ‘seed’ regions are behaviorally relevant.

### 4.6 Cognitive processes underlying ISFC correlations to behavior – perception versus memory

We asked how the movie content may affect the edge-behavior correlations and whether these correlations relate to the perception or the encoding of faces in memory. We calculated the IS-ETS correlation to behavior and examined how these resembled the global correlation obtained with the ISFC. We found that these similarities were highest during movie boundaries and hippocampus activations. In contrast, we did not find these similarities when faces appeared in the movie or during FFA activation.

Distributed functional connections of the hippocampus during memory encoding are thought to be necessary for the consolidation of perceptual segments in memory^69^. These connections might indicate the replay of perceived segments to promote encoding. They were also found with category-specific visual regions such as the FFA and PPA^70^, while the strength of these connections predicted subsequent memory^35^. Accordingly, memory for faces is considered essential for face recognition^71^, and it correlates with face perception abilities in typicals^72^ and CPs^73^ (but see^74^). Moreover, evidence suggests that memory deficits associated with faces play a significant role in the etiology of CP^75, 76^. Taking together, our results indicate that the observed cortical-wide interactions are related to, or at least more pronounced during, face encoding. To corroborate these results, future studies should include movie segments with and without faces and examine in which of them similar ISFC correlation to behavior may appear.

### 4.7 Limitations

We consider several limitations of the current study. First, the Benton Facial Recognition Test ^77^ used in this work was originally designed for diagnosing neurologically impaired individuals^78^ (but see^79^). More up-to-date standardized tests specifically designed to measure the normal range of face perception^72^ or memory^61^ might have better captured individual differences in our work. A second limitation relates to using a naturalistic movie in contrast to a well-controlled set of stimuli as typically used in perceptual neuroimaging experiments^80^. While using such stimuli to study brain-behavior relations has many advantages^81^, it is challenging to disentangle responses to different types of stimuli and cognitive processes, such as memory and perception in the current study. Moreover, we have no direct measure of subjects’ engagement level and focus of attention during the movie (but see ^82^).

### 4.8 Conclusions

Our work reveals how dynamics in cortical-wide functional interactions may support face recognition within the normal population. We found that these patterns are not limited to the classic ventral face patches but include regions that are implicated in attention, memory, and non-face selective visual processing. We report that these sets of connections co-fluctuate with activity within core face-selective areas, suggesting that the former might serve as hubs in this extended face network. Finally, we find evidence that individual differences observed in this network are related to the encoding, rather than the visual perception of faces. We propose that the methodological advancement made here can further explore brain-behavior relations in other cognitive domains.

## 5. Material and Methods

### 5.1 Participants

The data was obtained from the Cambridge Centre for Ageing and Neuroscience (Cam-CAN; ^21, 22^) dataset that includes functional and structural brain magnetic resonance imaging (MRI) along with demographics and face recognition scores. The Cam-CAN dataset includes 652 subjects (333 females, 322 males) aged 18-88 roughly uniformly distributed from Cam bridge City, UK. All participants provided informed consent, and the study was approved by the local ethics committee, Cambridgeshire 2 Research Ethics Committee (reference: 10/H0308/50). The data is freely available upon online access request https://camcan-archive.mrc-cbu.cam.ac.uk/dataaccess/. Additional information on the recruitment, eligibility criteria and demographics of the sample is available in the relevant publication^22^. Inclusion criteria were based on the availability of functional and structural imaging data, face recognition score, and successful completion of the preprocessing and quality control stages specified in the methods sections 2.3 and 2.4. The remaining subjects were assigned to groups according to their face recognition scores while keeping age distribution equal. A total of 517 subjects complied with these criteria and were included in all following analyses.

### 5.2 MRI acquisition

The data was acquired on 3T Siemens TIM Trio System, equipped with a 32-channel head coil. T1 structural images were acquired using a gradient echo (MPRAGE) sequence with TR=2250ms, TE =2.99ms; TI =900ms; flip angle =9°; FOV =256mm x 240mm x 192mm; voxel size =1mm isotropic. The movie fMRI session was acquired using a multi-echo EPI sequence with TR=2470ms, TE =9,4, 21.2, 33, 45, 57ms; flip angle =78°; FOV =192 x 192 x 142mm^3^, voxel size =3 × 3 × 3.44 mm^3^, slice thickness included 0.74mm gap and an acquisition time of 7min 57sec. In the fMRI session subjects were instructed to watch and listen to a movie.

### 5.3 MRI preprocessing

Preprocessing of structural and functional imaging was conducted using fMRIprep 20.0.6^83^ which is based on Nipype 1.4.2^84^. Parts of this section are adopted from the fMRIprep preprocessing report. The T1-weighted (T1w) image was corrected for intensity non-uniformity with N4BiasFieldCorrection^85^, distributed with ANTs 2.2.0^86^, and used as T1w-reference. The T1w-reference was then skull-stripped with a Nipype implementation of the antsBrainExtraction.sh workflow (from ANTs), using OASIS30ANTs as the target template. Brain tissue segmentation of cerebrospinal fluid (CSF), white-matter (WM) and gray-matter (GM) was performed on the brain-extracted T1w using fast (FSL 5.0.9^87^). Brain surfaces were reconstructed using recon-all (FreeSurfer 6.0.1^88^), and the brain mask estimated previously was refined with a custom variation of the method to reconcile ANTs-derived and FreeSurfer-derived segmentations of the cortical gray-matter of Mindboggle^89^. Volume-based spatial normalization to the MNI152NLin6Asym space^90^ was performed through nonlinear registration with antsRegistration (ANTs 2.2.0), using brain-extracted versions of both T1w reference and the T1w template.

The preprocessing of the movie fMRI data was as follows: First, a reference volume and its skull-stripped version were generated using a custom methodology of fMRIPrep. Susceptibility distortion correction (SDC) was omitted. The BOLD reference was then co-registered to the T1w reference using bbregister (FreeSurfer) which implements boundary-based registration^91^. Co-registration was configured with six degrees of freedom. Head-motion parameters with respect to the BOLD reference (transformation matrices, and six corresponding rotation and translation parameters) were estimated prior to any spatiotemporal filtering using mcflirt (FSL 5.0.9^92^). BOLD runs were slice-time corrected using 3dTshift from AFNI 20160207^93^. The BOLD time series including slice-timing correction were resampled onto their original, native space by applying the transforms to correct for head-motion. A T2* map was estimated from the preprocessed BOLD by fitting it to a monoexponential signal decay model with log-linear regression. For each voxel, the maximal number of echoes with reliable signal in that voxel were used to fit the model. The calculated T2* map was then used to optimally combine preprocessed BOLD across echoes following the method described in ^94^. The optimally combined time series was carried forward as the preprocessed BOLD. The BOLD time series were resampled into the MNI152NLin6Asym standard space. The following confound time series were regressed out: the global signal, framewise displacement (FD^95^), 6 motion estimates their derivatives and their squares, and 6 component-based noise correction from the CSF and WM (aCompCor^96^). Subjects with mean FD that was higher than 1.5 standard deviations from the group mean were excluded. This resulted in the exclusion of 39 subjects out of the total 567 subjects for whom face recognition measures and functional imaging were available.

### 5.4 Nodes and networks definition

Nodes were defined based on the Schaefer 400 cortical parcellation^97^ rendered on the MNI152NLin6Asym space. The ATL was defined according to the ‘AntTemp’ labels. The parahippocampal place area (PPA) was defined as the node with the maximal overlap to the “parahippocampal place” map (right hemisphere: rh_DefaultC_PHC_2, left hemisphere: lh_DefaultC_PHC_2) in Neuroquery^98^. The canonical resting-state networks were defined according to Yeo et al. 2011. The location of the bilateral posterior and medial fusiform face area (pFFA/mFFA) was determined using coordinates from previous Activation Likelihood Estimation (ALE) meta-analysis^99^ (rh pFFA: rh_VisCent_ExStr_3, rh mFFA: rh_VisCent_ExStr_1, lh pFFA: lh_VisCent_ExStr_3, lh mFFA: lh_VisCent_ExStr_1).

### 5.5 Face recognition scores

Face recognition ability was quantified using the accuracy score in the short form of the Benton Facial Recognition Test^77^ and a famous faces recognition test^22^. Benton’s test is a standardized test for assessing the recognition of unfamiliar faces. A target face was presented in each trial, followed by six test images. Subjects were required to identify one or more target faces among the six presented images despite lightning and head orientation changes. Scores were presented as proportions of correct responses out of 27 correct answers. In the famous faces test, participants were shown faces of famous people and unknown foils. They were instructed to identify the familiar faces and provide their names and occupation. In each of the 40 trials stating the name or occupation was sufficient to be considered a correct trial. The first unrotated principal component (PC) of the two measures was used as the main face recognition measure. Subjects with an accuracy score lower than 2.5 standard deviations from the mean in the combined test were excluded as outliers (11 excluded, 517 left).

### 5.6 Audiovisual stimulus

Participants watched a compelling black-and-white television drama by Alfred Hitchcock, “Bang! You’re Dead”^81^, edited from the original 30 min version to 8 min in a manner that preserved the main storyline^22^. To test which cognitive functions may relate to ISFC differences in face recognition, we extracted measures related to the visual perception or the encoding in memory of faces. We extracted measures related to the appearance of faces and their screen size on each movie frame (2Hz) using an ensemble of 4 pre-trained convolutional neural networks for face recognition/ classification^24, 100–103^. We computed the statistics for each TR (TR=2.47s) as the mean of the five frames analyzed during this TR. A movie depicting the face annotation is available online, along with the extracted measures and the code (https://github.com/GidLev/dynamic_faces_2022). Perceptual boundaries between movie segments, shown to precede the encoding of episodes or ‘chunks’ into memory^31^, were also taken from a previous study^23^. Each boundary between movie segments was rated by five to sixteen independent observers.

Raters watched the movie and identified points in time when “one event (meaningful unit) ended, and another began”^104^. All behavioral measures were normalized by a division by their maximum value.

### 5.7 Inter-subject functional connectivity (ISFC) analysis

ISFC is a computational method designed to measure the extent of synchrony between the brains of different subjects driven by a common stimulus^19^. It can be measured between homologous brain regions (e.g., same voxel or brain region) across subjects using inter-subject correlation (ISC) or between different brain regions (ISFC). We computed ISFC values as follows: the first and last five TRs (∼12 s) were excluded in all subjects to avoid onset and offset effects^18^. Then, the Pearson correlations between all possible node pairs were computed between each subject and all other members of its group (see 2.7 below for details). Finally, the correlation values were Fisher to z transformed. We used the ISFC standalone implementation from https://github.com/snastase/isc-tutorial. All glass and surface brain plots were done using Nilearn^105^.

### 5.8 Behavioral groups assignment

As in previous studies using ISFC, we measured shared neural responses between a subject and all other members of its group. However, mixing individuals with heterogonies face recognition abilities within the same group could mask their behavioral-relevant shared neuronal response. To overcome this limitation, subjects were divided into equal-sized groups according to their face recognition score while keeping their age distribution equal. Specifically, we divided subjects into bins according to their age (18-28,28-38,…,78-88). Then, in each age bin, subjects were assigned into *i* equal behavioral groups according to their face recognition score. We conducted all following analyses with *i*=6 (n=517) and additionally replicated the results using different choices of *i* (i=5,7; see supplementary information [SI] 9.1).

### 5.9 Inter-subject node and edge time series (IS-NTS/ IS-ETS) analysis

To explore temporal dynamics in the face processing network, we decomposed connectivity to its edge time series^20^. Here, instead of within-subjects, this was done across subjects to highlight their stimulus-dependent shared responses. This novel method, we named the inter-subject node or edge time series (IS-N/ETS), quantifies the extent of inter-subject connectivity co-fluctuation (“synchrony”). The IS-NTS is computed as the element-wise multiplications of the z-scored signals of homologous brain node across subjects (e.g., same voxel or brain region). To differentiate between co-activation and co-deactivation, we multiply this time series by the sign of the node’s signal. The IS-NTS can be viewed as an approximation of stimulus-dependent shared activations. The IS-ETS can be similarly derived between distinct brain regions/voxels by avoiding the multiplication by the sign step. We refer to SI section 9.2 for a numerical simulation demonstrating the advantage of the two measures in capturing stimulus-dependent shared co-activation/co-deactivation and connectivity co-fluctuations.

### 5.10 Edge seed correlation

To examine whether brain-wide co-fluctuations (‘targets’) may covary with the activity of a given brain region (‘seed’), we developed the ’edge seed correlation’ method. Using our novel IS-NTS/ETS method (see 2.8), this can be conducted in a stimulus-driven manner by testing the correlation between the IS-NTS of a specific brain region and all possible IS-ETS across the brain. Edges’ significance was determined by comparing their values to a null distribution (k=10,000) created by applying phase randomization on the IS-NTS^106^. Randomization was done by Fourier transforming the IS-NTS, randomizing the phase of each component, and applying the inverse Fourier transformation. From computational time considerations, in cases were a stricter threshold than 1/k (p<0.0001) was applied, we used the p-value computed from the number of standard deviations from the mean of the null distribution. The edge seed correlation could also be applied by correlating regions’ signal and brain-wide edge-time series. In the relevant analyses (see 3.3 and SI 9.3), we examined whether the IS edge seed correlation may increase the observed statistical power over the non-IS variate of the test. We computed effect size across the two variants of the test by thresholding both correlation matrices (|r| > 1^st^ percentile). Then, we calculated its distance from the mean of the null distribution for each edge divided by the standard deviation. We report the mean power resulting from each test and their difference using a paired t-test. We share a Python code and interactive notebook demonstrating the IS-N/ETS derivation and the IS edge seed correlation method (https://github.com/GidLev/dynamic_faces_2022).

### 5.11 Statistical analysis

In all analyses, Spearman’s correlation coefficients are denoted by the letter ρ and Pearson’s by the letter r. The similarity between edge maps was quantified using the Dice similarity coefficient. For non-binary maps, dice similarity was measured after map thresholding (1st percentile). Brain map decoding was done by testing its Dice similarity compared to a dictionary of 128 cognitive brain maps (Cogspaces; (Mensch et al., 2021) and reporting the map with the highest similarity. For visualization of the selected cognitive map, a word cloud representation was derived using the Neurosynth image decoder (Yarkoni et al., 2011). The hemodynamic response function (see Results 3.3) was defined as in https://bic-berkeley.github.io/psych-214-fall-2016. P-values for the correlation between behavioral predictors in Result section 3.4 were determined using a phase-shuffling procedure (see 2.9). Neural events were computed by taking the mean IS-ETS across subjects, then the root-mean-square (RMS) across all edges for each time-point^20^.

## 6. Data and code availability

The unprocessed data is openly available upon online access request https://camcan-archive.mrc-cbu.cam.ac.uk/dataaccess/. The code for reproducing the IS-N/ETS and the IS edge seed correlation methods along with the face predictors and the code that generates them, is available online (https://github.com/GidLev/dynamic_faces_2022).

## Acknowledgments

Data collection and sharing for this project were provided by the Cambridge Centre for Ageing and Neuroscience (CamCAN).

Corresponding author E-mail address: (G. Levakov).

