## Supplementary material for "Fine-scale dynamics of functional connectivity in the face processing network during movie watching": SI

### 10. Supplementary information (SI)

#### *10.1 ISFC and its relation to face recognition behavioral performance – different number of behavioral groups*

Section 3.1 outlined the correlation between participants' ISFC values and face recognition behavioral group affiliation ($i$=6) for all possible edges. Here we describe the results when using an alternative number of behavioral groups.

$i$=5: We found that edges that negatively correlated to recognition scores were mostly between visual and attentional dorsal (9.09%) or frontal default (72.73%) network nodes. The dorsal attentional nodes are part of the bilateral PPC, and the frontal default nodes mostly include the medial prefrontal cortex (mPFC) and the inferior frontal gyrus (IFG). Only three positively correlated edges were found significant, hence we additionally examined edges using a more lenient threshold ($p<0.001$). This revealed that the nodes with the highest degree for the negative configuration were the rh. lateral occipital cortex (LOC) (degree=14). The lh. ATL was the highest degree node for the negative configuration (degree=10).

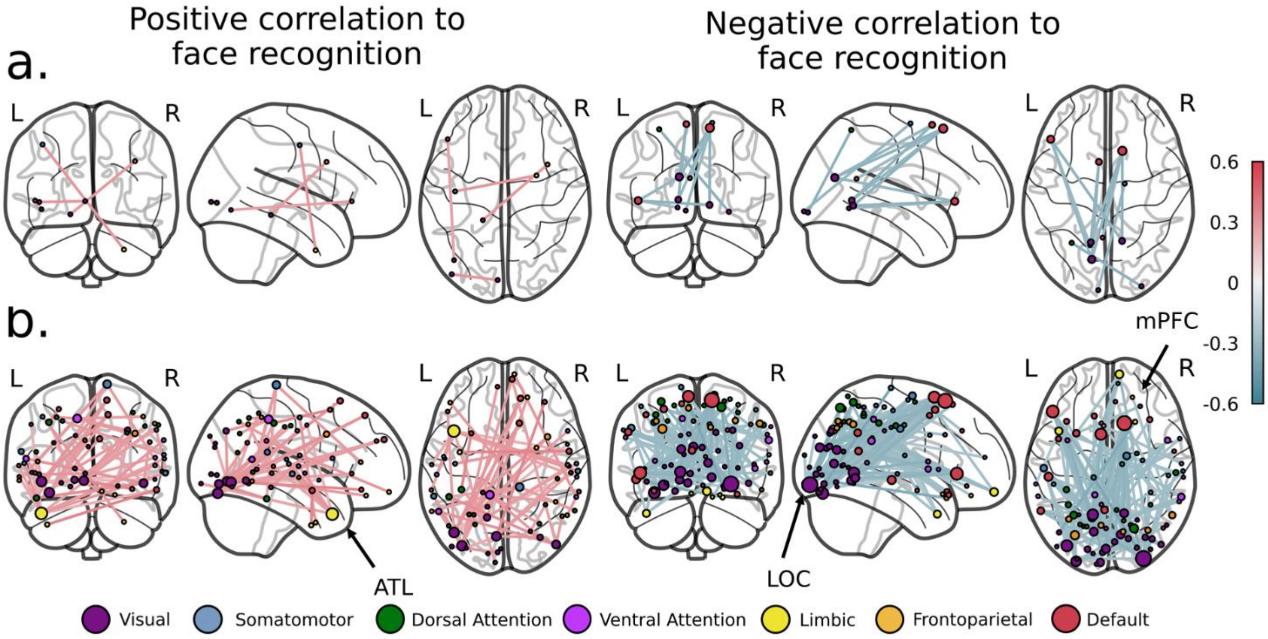


**SI Fig. 1 - Correlation of ISFC values and face recognition ability for five behavioral groups (*i*=5). For the full figure legend see Fig. 1.**

<u>*i*=7</u>: We found that edges that negatively correlated to recognition scores were mostly between visual and attentional dorsal (7.69%) or frontal default (61.54%) network nodes. The dorsal attentional nodes are part of the bilateral PPC, and the frontal default nodes mostly include the medial prefrontal cortex (mPFC) and the inferior frontal gyrus (IFG). Only three positively correlated edges were found significant, hence we additionally examined edges using a more lenient threshold ($p<0.001$). This revealed that the nodes with the highest degree for the negative configuration were the rh. lateral occipital cortex (LOC) and the mPFC (both degree=11). The lh. ATL was the highest degree node for the negative configuration (degree=17).

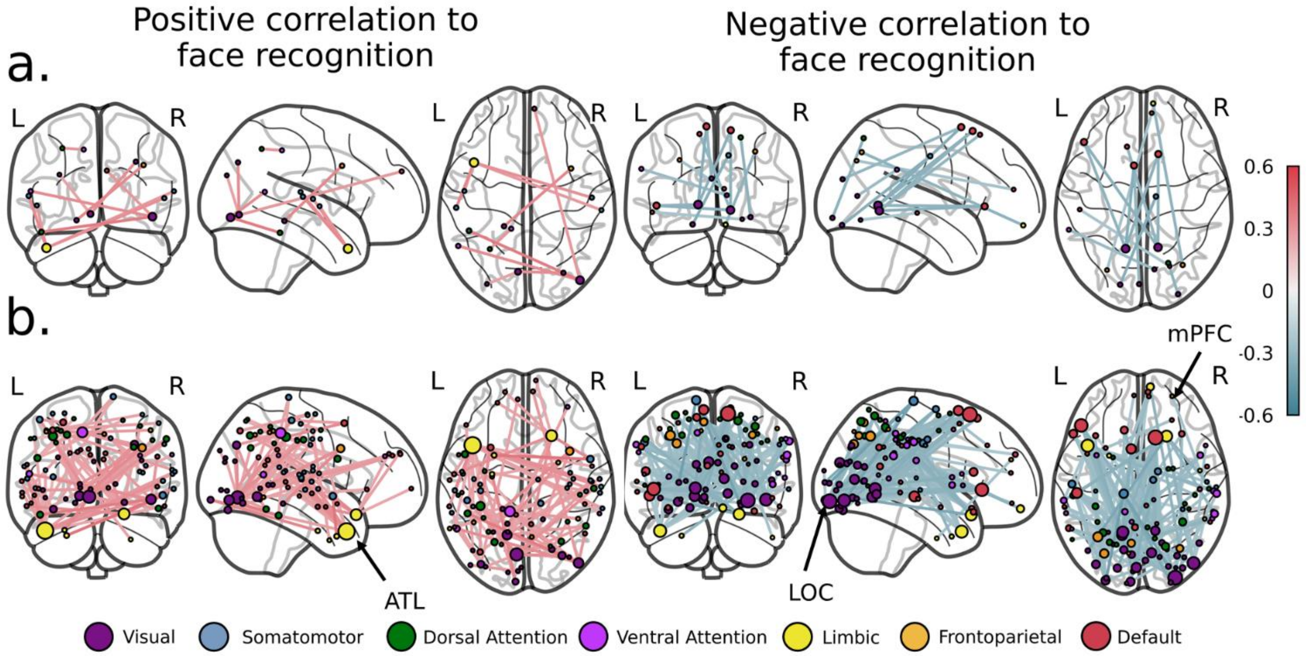


**SI Fig. 2 - Correlation of ISFC values and face recognition ability for five behavioral groups (*i*=7). For the full figure legend see Fig. 1.**

#### *10.2 Evaluating the relation of IS-NTS/ETS to stimulus-locked activity/connectivity using numerical simulation*

Using a numerical simulation, we tested the extent that our IS node/edge time series (*IS-NTS/ETS*) methods capture stimulus-locked activity and connectivity co-fluctuations. We simulated the neural activity of two brain regions, a and b, during a typical fMRI experiment. The experiment included 40 subjects that viewed a shared stimulus with a length of 300 TRs. The activity of each region at time *t* was described as:

$$x_t = \alpha_t \cdot c_t + (1 - \alpha_t) \cdot n_t$$

Such that *c* was the subjects' shared response, *n* was the idiosyncratic response/noise, and *α* was the weights that balanced the two. All three are vectors of length equal the number of TRs and randomly set with a uniform distribution [*c,n ~U(-1,1); α ~U(0,1)*]. *c* and *α* were identical for all subjects and regions, and $n$ were unique for each subject and region. We computed the correlation of region a's weighted shared signal ($\alpha \cdot c$) with the IS-NTS and the original signal ($x$). We found a significantly higher correlation ($t=85.219$, $p=6.23e-46$) between the weighted shared signal and the IS-NTS ($r=0.906 \pm 0.007$) compared to the original signal ($r=0.697 \pm 0.018$). Next, as $c$ was common to both regions a and b, it effectively drove their shared co-fluctuations. Hence, we compared the amplitude of the shared co-fluctuations ($|\alpha \cdot c|$) to both the edge-time series and the IS-ETS between regions a and b. We found a significantly higher correlation ($t=41.074$, $p=1.03e-33$) between the weighted shared signal and the IS-NTS ($r=0.853 \pm 0.009$) compared to the original signal ($r=0.680 \pm 0.033$; SI Fig. 3).

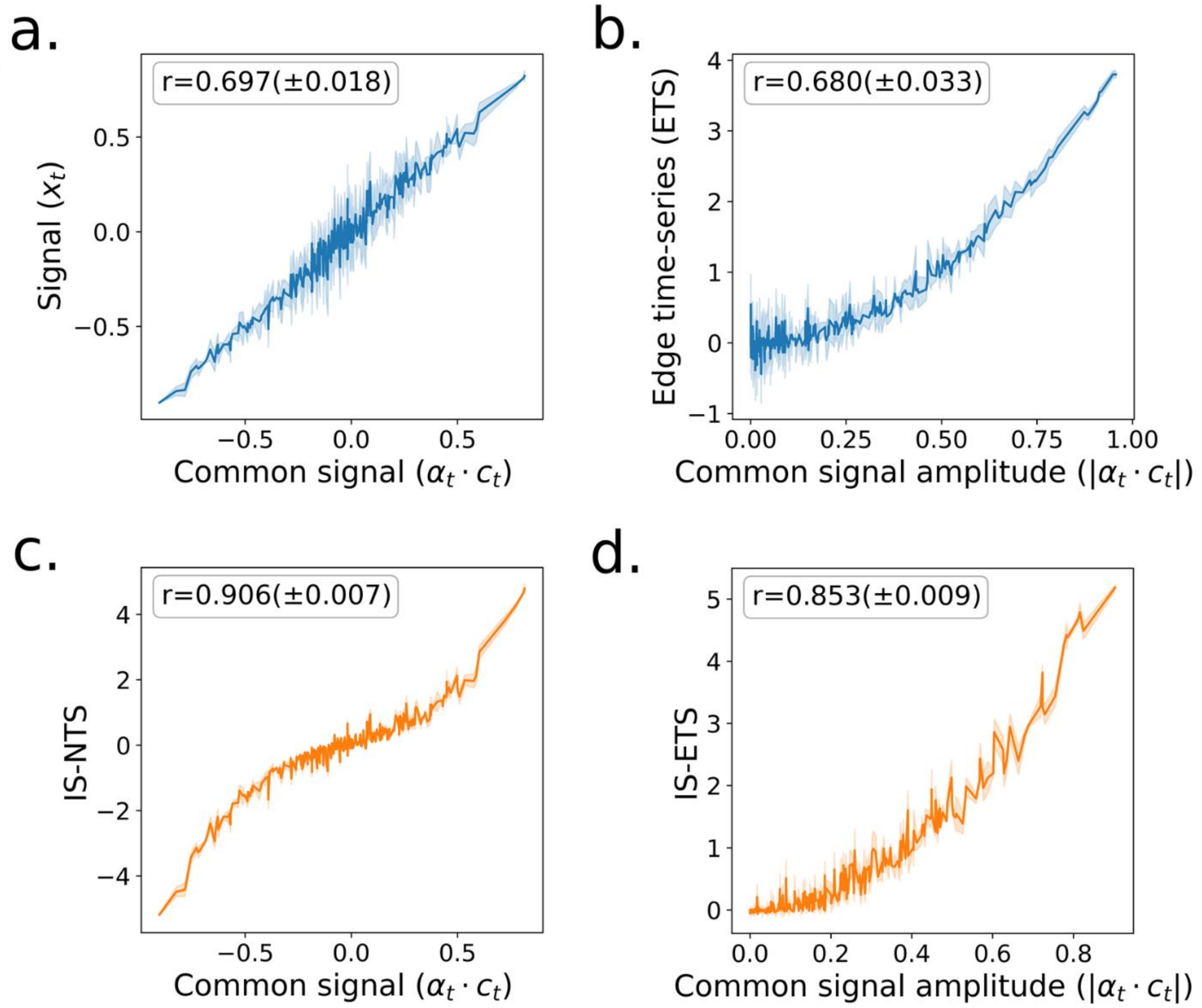


$$x_t = \alpha_t \cdot c_t + (1 - \alpha_t) \cdot n_t$$

**SI Fig. 3 - Correlation of the IS-NTS/ETS to stimulus-locked activity/connectivity using numerical simulation. The left panels depict the correlation of the weighted shared signal (α · c) of region a with the IS-NTS (a) and the original signal (c). The right panel depicts the correlation of the amplitude of the shared co-fluctuations (|α · c|) to both the edge-time series (b) and the IS-ETS (c) between regions a and b. The simulated signal is defined using the equation on the bottom such that *c* was the subjects' shared response, *n* was the idiosyncratic response/noise, and *α* was the weights that balance the two.**

#### *10.3 Validation of edge-seed correlation method within the frontal insula*

We validate the edge-seed correlation method by replicating findings within the frontal insula, previously shown to drive the switching between the default mode and the frontoparietal networks [26,27]. We hypothesized that given the role of the frontal insula in switching between the two systems, seeding it would reveal positive correlations to edges connecting frontoparietal nodes and negative correlations to edges connecting default mode nodes. The network affiliation of nodes was defined according to Yeo's 7-networks definition [25].

As expected, seeding the rh. frontal insula revealed a positive correlation to edges connecting the frontoparietal network nodes (56.97% of significant edges) and a negative correlation to edges connecting the default network nodes (34.74% of significant edges; SI Fig. 4). Importantly, most significant edges were not directly connected to the seeded node or its homologous node in the left hemisphere (88.05%).

The edge seed correlation method could also be applied within (intra) subjects by correlating node time series with the edge time series of all possible edges. We computed the intra-subject edge-seed correlation and compared it to its inter-subject. Effect size across the two test variants was defined by thresholding both correlation matrices ($|r| > 1^{st}$ percentile) and taking the resulting mean z-scores. Computing the edge seed correlation within, instead of across subjects, produced a significantly smaller effect size (Cohen's d: Inter-subject=4.48, Intra-subject=2.63; $t(1598)=120.292$, $p<2.2e-16$), suggesting an advantage of applying this method across subjects.

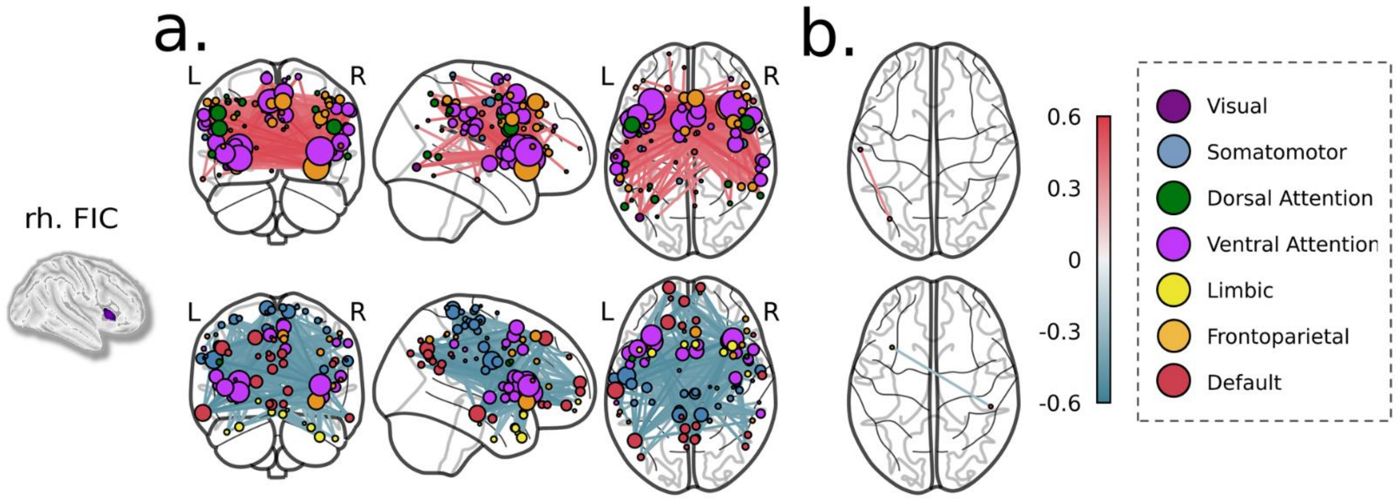

SI Fig. 4 – IS edge seed correlation of the rh. frontal insula. IS edge seed correlation is defined as the IS correlation between the node time series (NTS) of a specific region and the time series of all possible edges (ETS). (b) A glass brain projection of edges correlated to the rh. frontal insula (p<1.0e-05). As expected, IS-NTS of the frontal insula was positively correlated to edges connecting the frontoparietal network (56.97%) and negatively to edges connecting the default mode network (34.74%). Only one edge survived the same threshold when repeating the test intra instead of inter subjects (right side). The color of the nodes indicates their affiliation to one of the seven canonical resting-state networks (Yeo et al., 2011). Nodes' size is proportional to their strength, i.e. the sum of their absolute edge weights. Edges are projected onto coronal (left), sagittal (middle), and horizontal (right) views.

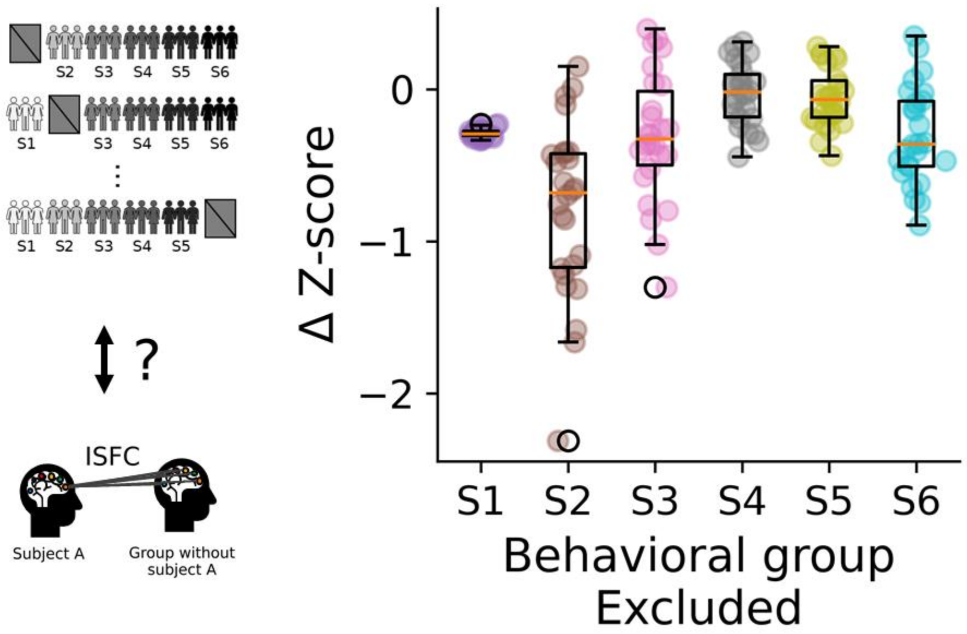


SI Fig. 5 – Box plot depicting the effect of removing one of the six behavioral groups on the relation to behavior. Each point represents an edge that was found significant using all groups. The x-axis represents the excluded group and the y-axis the change in the edge z-score such that an increased correlation to behavior is positive and vice versa. The z-score of each edge was computed as its difference from the mean divided by the standard deviation of the null distribution. The effect of group exclusion on each significant edge was computed as [after exclusion] minus [before exclusion] for positively correlated edges and vise-versa for negatively correlated edges.

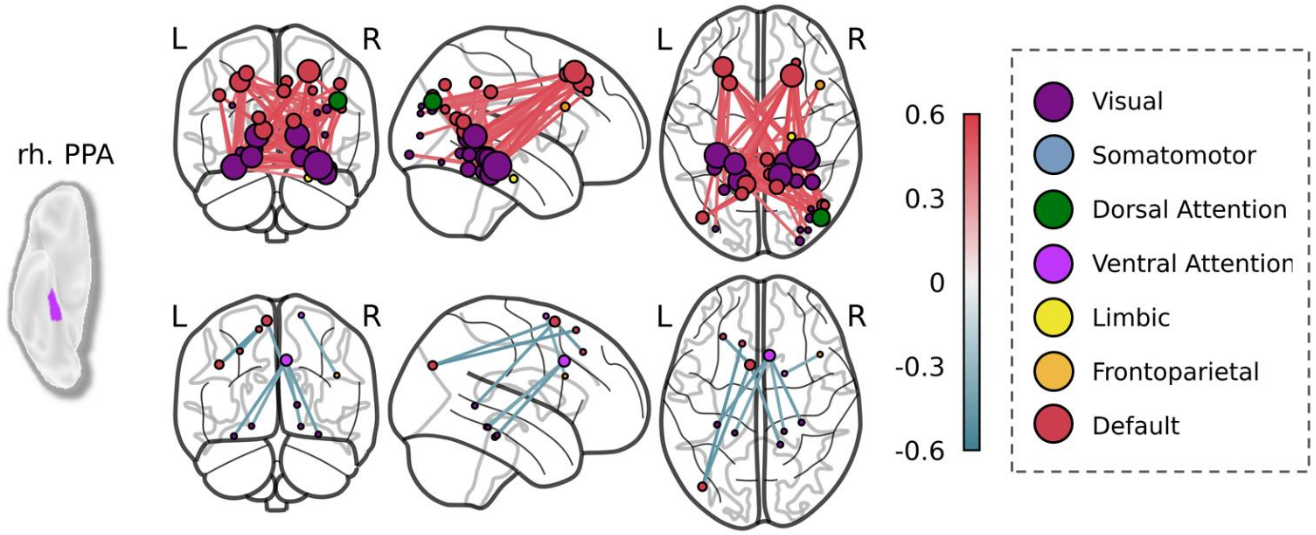


**SI Fig. 6 – IS edge seed correlation of the rh. parahippocampal place area (PPA). IS edge seed correlation is defined as the IS correlation between the node time series (NTS) of a specific region and the time series of all possible edges (ETS). (b) A glass brain projection of edges correlated to the rh. PPA (p<1.0e-05). The color of the nodes indicates their affiliation to one of the seven canonical resting-state networks (Yeo et al., 2011). Nodes' size is proportional to their strength, i.e. the sum of their absolute edge weights. Edges are projected onto coronal (left), sagittal (middle), and horizontal (right) views.**

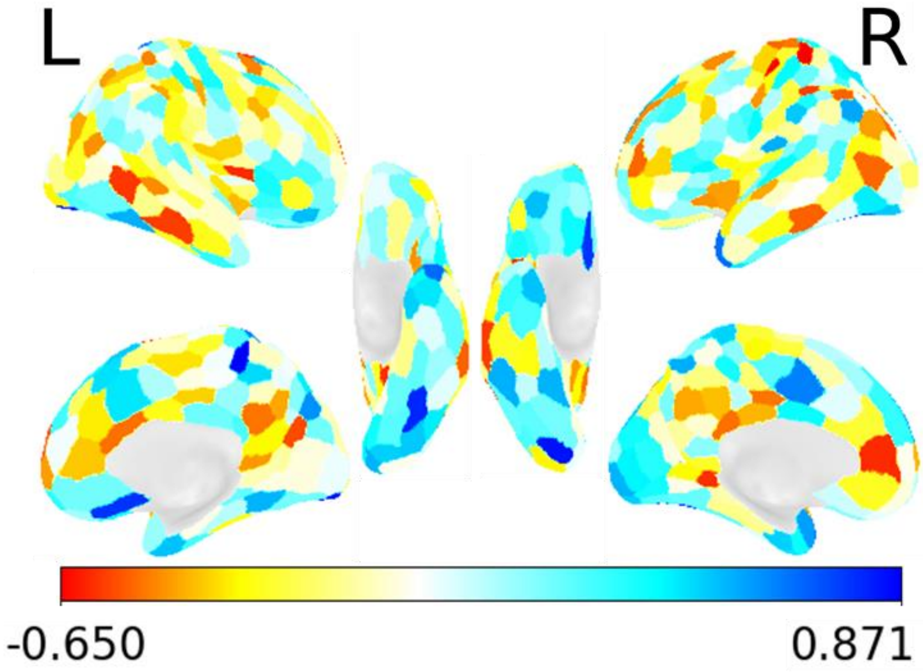

**SI Fig. 7 - Correlation of groups' regional activation differences to behavioral differences. Surface plot depicting the correlation of groups' regional activation differences to groups' behavioral differences.**

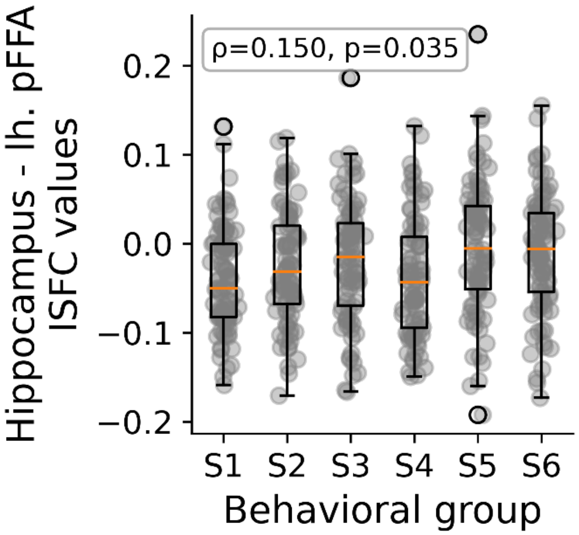


**SI Fig. 8 - Hippocampal-FFA ISFC correlation with and behavior. Box plot depicting ISFC for the bilateral hippocampus and the lh. pFFA for each behavioral group. Each dot represents a single subject. We found a positive Spearman correlation between behavioral group affiliation and ISFC values (ρ=0.150, p=0.035).**
